# The attractor landscape of duplicated networks

**DOI:** 10.1101/2023.01.30.526352

**Authors:** Enrico Borriello

## Abstract

We study the effect that network duplication has on the topology of the state space of dynamical Boolean networks with thresholds. We show that the problem of finding the attractor dynamics is just as hard as finding the attractors of the unduplicated network. We also show that a reverse algorithm –normally not computationally advantageous in determining the basins of attraction– can now exploit the symmetry of the system and its computational complexity does not scales exponentially anymore with the size of the network. Lastly, we show that when a chain of network duplication events is considered, only the first events change the nature of the attractors, while successive events only affect/reinforce the basins of attraction.

---

Since Kauffman’s initial proposal [1], dynamical Boolean networks have been frequently adopted in systems biology as simple, mechanistic models of gene regulation. In this framework, nodes represent genes/operons, and their associated dynamical variables are Boolean approximation of their expression. Therefore, they only possess two possible states: either active or inactive. Links represent gene regulation, and the attractor states of the dynamics –fix-points and/or cycles– represent cell types/phenotypes [2, 3, 4, 5].

In this manuscript we explore the topology of the state space of a duplicated network, with a special emphasis on the computational simplification that the symmetry of the system imposes on the nature of the attractors The interpretation of Boolean networks as simplified models for gene regulatory networks (GRNs) allows to reinterpret this manuscript as an idealized attempt at exploring the connection between the phenotypes of a genome, and those of the corresponding tetraploid [6, 7]. We will mostly focus on the mathematical aspect of the problem, but we will reinterpret some of our results in the light of systems biology in our discussion section. In what follows we will always assume complete cross-talk between the original network and its replica (perfect polyploidy) [8], and leave the generalizations needed to include aneuploidy (e.g. chromosome loss [9]) for a follow-up of this manuscript.

## Methods

### Boolean networks with thresholds

The state of a Boolean network at time *t* is defined by the state of its *n* nodes, *x*_*i*_(*t*), with *i* = 1, …, *n*. The Boolean nature of the model means that there are only two possible states for node *i*: ON, which corresponds to *x*_*i*_(*t*) = 1, or OFF, with *x*_*i*_(*t*) = 0. In these manuscript we will consider the case of (synchronous) Boolean networks with thresholds. In these models, the time evolution of the dynamic variables *x*_1_, …, *x*_*n*_ is given by

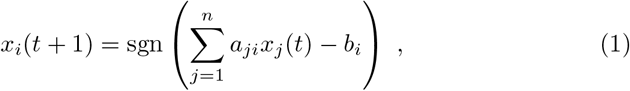

where *a*_*ji*_ is the relative weight of the regulatory signal from node *j* to node *i* (activation when *a*_*ji*_ is positive, inhibition otherwise), and *b*_*i*_ is the activation threshold of node *i*. The threshold function sgn(*x*) is just the unitary step function: sgn(*x*) = 0 if *x* ≤ 0 but sgn(*x*) = 1 if *x >* 0. In a more compact notation, we can write

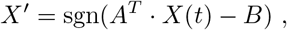

where we have introduced the *adjacency matrix A* = (*a*_*ij*_), and the columns *B* = (*b*_*i*_) and *X* = (*x*_*i*_).

Examples of the application of this class of models to gene regulation can be found in [10, 11].

### Network duplication

Given a Boolean network *N*_1_ with *n* nodes, and *n* small enough as for us to assume its dynamics can be solved, we want to study the dynamics of the 2*n*-node network *N*_2_ obtained by duplicating each node of *N*_1_.

We will assume that the dynamics of *N*_1_ is governed by the following set of Boolean equations

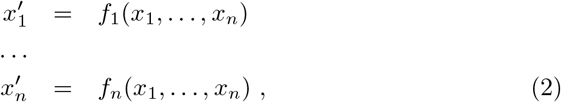

where *x*_*i*_ is the value (0 or 1) of node *i* at time *t*, and 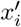 is its value at time *t* + 1. The updating rules of *N*_2_ will have a similar form

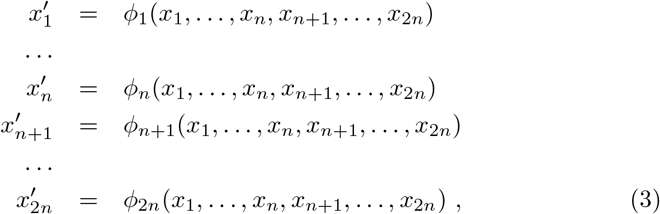

where the *ϕ*_*i*_ functions are new functions that, in general we do not know. Our task to establish assumptions about the functional form of these *ϕ*_*i*_ functions, based on our knowledge of the functions *f*_*i*_. Following [5], we can build the duplicated network via the inclusion of one node replica at a time, starting from node *n* + 1, assumed to be a perfect replica of node 1, and then proceeding in a similar way with nodes *n* + 2, …, 2*n*. Let us then consider the requirements we impose on *ϕ*_*i*_(*x*_1_, …, *x*_*n*_, *x*_*n*+1_), after we include node *n* + 1:

1. The observation that node *n* + 1 is a *perfect replica* of node 1 translates into the assumption that

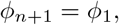

i.e. we are imposing that node replicas obey the same updating rule.
2. We require that a new node does not affect the system unless it is expressed. Therefore,

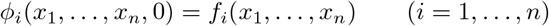
3. Our last assumption establishes the effect that node replicas have on the remaining nodes, and we impose that, to nodes 2, …, *n*, node 1 and *n* + 1 are *indistinguishable*:

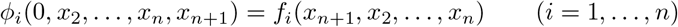

Using the previous conditions we can deduce the values that the new functions *ϕ*_*i*_ take for any state of *N*_2_, except states where *x*_*i*_ and *x*_*n*+*i*_ are both 1. These represent genuinely new scenarios, and correspond to output that cannot be deduced from of an equivalent configuration of *N*_1_. Notice that our discussion so far hold for generic Boolean networks. But, at least in Boolean networks with threshold we can circumvent this problem because of the prescription that generates the updating rule of a node in terms of the state of the network, eq. (1). Building the functions *ϕ*_*i*_ in an analogous way we are lead to impose *a*_*j*+*n,i*_ = *a*_*ji*_ and *b*_*i*+*n*_ = *b*_*i*_ for the previous three conditions to be satisfied.

Now that we know how to build *N*_2_ starting from *N*_1_, let us focus on the problem of solving its dynamics. Even when the dynamics of *N*_1_ can be solved in an exact way, this might still not be the case for *N*_2_, given its larger size. For example, solving the dynamics of a Boolean network with 10 nodes translates into the study of the topology of a directed network (the state space) with about just 1k nodes. But the state space of a dynamical network with 20 nodes already contains about 1M states. In the next subsections we will to show how to circumvent this problem enforcing the symmetry of *N*_2_.

### Symmetry

The fact that *ϕ*_*n*+*i*_ = *ϕ*_*i*_ is no guarantee, in general, that *x*_*n*+*i*_(*t*+1) will be equal to *x*_*i*_(*t* + 1). But this is the case with our parametrization of the network^1^, as our dynamical rules (1) update *x*_*n*+*i*_ and *x*_*i*_ to the same value. The values of the two nodes can still differ at *t* = 0, but they will get synchronized after a single time step. This observation implies a huge simplification of the dynamics of *N*_2_, and makes the problem of finding its attractors solvable anytime the same problem can be solved for *N*_1_. More specifically, we want to prove the following

#### Preposition 1

We can find the attractors of *N*_2_ by solving the dynamics of the virtual *n*-node network obtained by *N*_1_ after the replacement of *A* with 2*A* or, equivalently, *B* with *B/*2.

**Proof**

To prove it, let us first call *S*_1_ the configuration space of *N*_1_, and *S*_2_ the configuration space of *N*_2_. The number of elements in *S*_1_ is ord(*S*_1_) = 2^*n*^, while ord(*S*_2_) = 2^2*n*^ = (ord(*S*_1_))^2^. Therefore, when the network gets duplicated, the number of its dynamical states gets squared. Let us now define the set

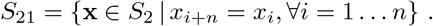

*S*_21_ is isomorphic to *S*_1_. But, at this level, this correspondence is just a set isomorphism: A one-to-one mapping between the elements of the two sets. We can rephrase the observation that *x*_*n*+*i*_(*t* + 1) = *x*_*i*_(*t* + 1) by saying that, after the first time step, all the states in *S*_2_ transition toward states in *S*_21_ (see figure 1). Equivalently, we can say that states in the complement of *S*_21_, let us call it 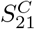, have no *pre-images*. Adopting the very romantic terminology in use in network dynamics, we will refer to states with this property as *Garden of Eden* (GoE) states [12]. Their number parametrizes the degree of irreversibility of a finite, deterministic system. Besides, they play a peculiar role in the dynamics of the system under study: Every additional dynamical state the network acquires after duplication (the states in 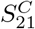) is just a GoE state. This also means that all the attractors of *N*_2_ are in *S*_21_.

**Figure 1:**
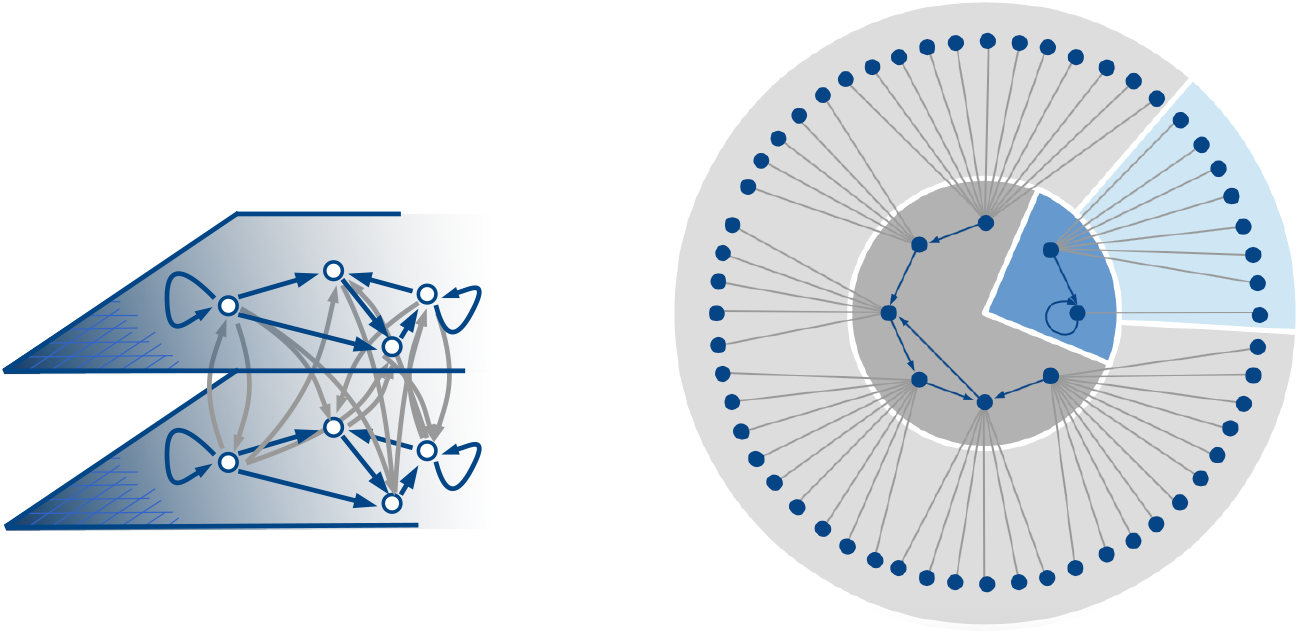
(Left) Topology of a duplicated network. (Right) Topology of the attractor landscape of a duplicated network with just 3 initial nodes, and 64 final states. 56 of these states are just GoE states evolving toward *S*_21_ states after a time step.

The problem is then reduced to solving the dynamics of a virtual network with just *n* nodes (replica nodes do not convey additional degrees of freedom), whose configuration space is now *S*_21_. This problem is trivial in the case of Boolean networks with thresholds, like the ones we are considering here. As we saw, the functions *ϕ*_*i*_ in eq. (3) are of the form:

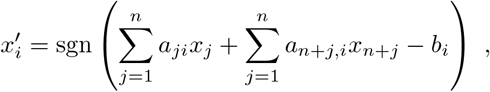

with *a*_*n*+*j,i*_ = *a*_*ji*_. Furthermore, after the first time step (when the dynamics is entirely contained in *S*_21_) *x*_*n*+*j*_ = *x*_*j*_, and the previous updating rule simply becomes

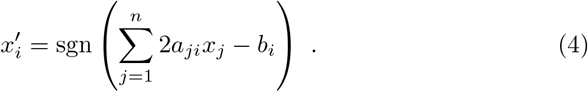

Therefore, the attractors of *N*_2_ can be deduced from the dynamic of the virtual *n*-node network obtained by *N*_1_ after the adjacency matrix *A* is replaced with 2*A* or, equivalently, after the thresholds *B* are replaced with *B/*2.

If *X* is an attractor state of this virtual network, then (*X, X*) is an attractor state of *N*_2_. It should be clear from the previous proof that finding the attractors of *N*_2_ is not more cumbersome than finding the attractors of *N*_1_. What the previous method is not proving are the new basins of attraction. The next subsection is devoted to addressing this problem.

### Reverse algorithm

Let us refer to a generic Boolean network with *n* nodes and updating rules like in (2). We will call *S*_*i,b*_ the subset of the configuration space *S* containing all the states satisfying the non-linear equation *f*_*i*_(*x*_1_, …, *x*_*n*_) = *b*, where *b* = 0, 1. Therefore, given a state *X* = (*x*_1_, …, *x*_*n*_), the set of its pre-images is

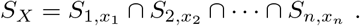

Finding the 2*n* sets *S*_*i,b*_ is equivalent to finding the pre-images of each state in *S*. A trivial algorithm for finding *S*_*i,b*_ would evaluate *f*_*i*_(*x*_1_, …, *x*_*n*_) for all (*x*_1_, …, *x*_*n*_) ∈ *S*, and select all the states for which *f*_*i*_(*x*_1_, …, *x*_*n*_) = *b*. This approach would require 2*n*× 2^*n*^ operations, as opposed to the *n* ×2^*n*^ operations we would need to determine the forward evolution of the network, and to draw the attractor landscape. In general, using a reverse algorithm is not a better way of solving the dynamics of a network [13]. But we can show that, for our duplicated network, reverse algorithms are especially efficient. First, though, we need to *compress* the sets *S*_*i,b*_ in a convenient way.

If we call *P*_*i,b*_ the set of sets of parametric equations defining the elements of *S*_*i,b*_, then the set of sets of equations defining the elements of *S*_*X*_ will be

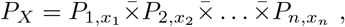

where 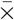 denotes an *unordered* and *repetition-free* Cartesian product of sets, from which sets containing *inconsistent equations have been removed*.

#### Example 1

To clarify our notations, let us refer to a very simple example, and consider a network with just 3 nodes and updating rules

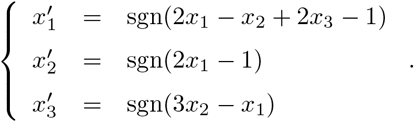

Let us find the pre-images of state *X* = (1, 1, 1). From the truth table of *f*_1_(*x*_1_, *x*_2_, *x*_3_) = sgn(2*x*_1_ − *x*_2_ + 2*x*_3_ − 1) we can easily obtain

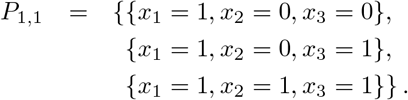

Analogously, we can derive

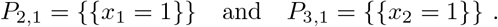

Therefore,

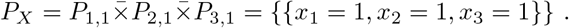

In the previous product, sets of equations containing *x*_1_ = 1 twice have been simplified. Sets containing both the inconsistent equations *x*_2_ = 0 *and x*_2_ = 1 have been removed, etc. From *P*_*X*_ we can now deduce *S*_*X*_ = {(1, 1, 1)}, i.e. *X* is its own only pre-image.

Notice that, while *P*_1,1_ and *S*_1,1_ = {(1, 0, 0), (1, 0, 1), (1, 1, 1)} contain the same number of elements, *P*_2,1_ and *P*_3,1_ each compresses a set with four states using just one parametric equation.

It is easy to show that two more attractors exist, the isolated fix-point (0, 0, 0), and the cycle {(0, 0, 1), (1, 0, 0), (1, 1, 0), (0, 1, 1)}.

Even if the sets *S*_*i,b*_ were all know, it is still computationally convenient to use the set of equations *P*_*i,b*_. While the order of the *S*_*i,b*_ sets grows exponentially with the number of nodes *n*, each elements in *P*_*i,b*_ contains at most *k*_*i*_ equations, where *k*_*i*_ is the in-degree of node *i*, and there are at most 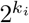 equations in *P*_*i,b*_. The complexity of the problem now scales exponentially with respect to the connectivity of the network, not its size. *S*_*i,b*_ and *P*_*i,b*_ convey the same amount of information, but this information is compressed in the latter sets. Besides, we can perform the main operation over the *S*_*i,b*_ sets –namely intersection– using the sets *P*_*i,b*_, and without the need of decompressing them.

This simplification holds only as long as we refer to the problem of finding the pre-images of a single node. In finding the basins of attraction, we would still need to repeat this process for almost every node in the network, unless some symmetry argument can be used to simplify the calculation.

In general, given a reverse algorithm for finding the pre-images, the search for the basins of attraction proceeds like his: We first start from a state, let say *X*_0_, and we follow its forward evolution until we reach an attractor state. At that point, we find the pre-images of all the states in that attractor, the pre-images of these newly found states, and so on, until the search fails to find new states. At that point we know the the basin of attraction *X*_0_ belongs to. After that, we choose a new state outside of that basin, and repeat this process until we determine the size of the new basin it belongs to. We keep repeating this process until we saturate the entire configuration space *S*. Next we want to prove that this process is enormously simplified in the case of a Boolean network with thresholds *N*_2_ obtained by duplicating all the nodes of an initial network *N*_1_.

#### Preposition 2

The problems of finding the basins of attractions of networks *N*_1_ and *N*_2_ through a reverse algorithm possess the same computational complexity.

**Proof:** As the equations for node *i* and node *n* + *i* (*i* = 1, …, *n*) are the same, it follows that *P*_*i*+*n,b*_ = *P*_*i,b*_. The new sets *P*_*i,b*_ depend now on the 2*n* variables *x*_1_, …, *x*_*n*_, *x*_*n*+1_ …, *x*_2*n*_, but we need to evaluate only half of the sets. Besides, as we need to find the pre-images of states in *S*_21_, for which *x*_*n*+*i*_ = *x*_*i*_, ∀*n*, the product

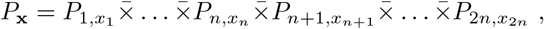

simply becomes

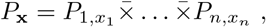

because 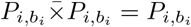.

But the main simplification comes from the fact that, outside of *S*_21_, there are only GoE states. Therefore, we do not need to step back any further once we exit *S*_21_. This is how the exponential catastrophe is prevent.

#### Example 2

With *N*_1_ as in example 1, let us consider the network *N*_2_ obtained by duplicating *N*_1_. In finding the attractors, we only need to consider the dynamics of an equivalent network with 3 nodes, after we multiply its adjacency matrix by 2:

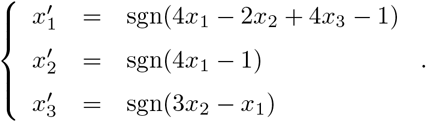

We can then deduce the new *P* -sets:

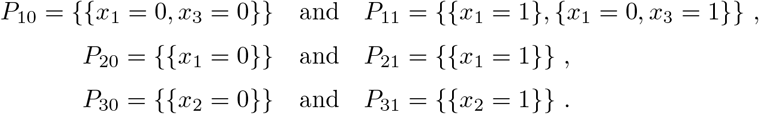

Taking the 8 possible 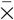 products of these sets is computationally equivalent to directly solving the dynamics of the network before duplication, and leads to just two solutions: (0, 0, 0) and (1, 1, 1), meaning that the duplicated network has attractors *X*_0_ = (0, 0, 0, 0, 0, 0) and *X*_1_ = (1, 1, 1, 1, 1, 1). The basin size of each attractor is then calculated reapplying the reverse algorithm only as long as the pre-images are still in *S*_21_, and remembering that *P*_*i*+*n,b*_ = *P*_*i,b*_ ∀*i* = 1 …, *n* (with *n* = 3 in this case). Starting with *X*_0_:

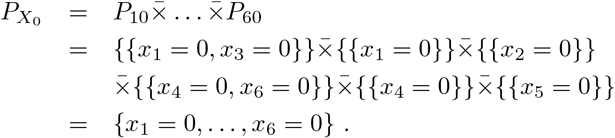

In this simple case, *X*_0_ is an isolated fix-point, while the remaining 63 states form the basin of *X*_1_.

As a side note, the formalism developed here can also be applied to the search of the GoE states of the unduplicated network *N*_1_, and for more general classes of dynamical Boolean networks than just the threshold networks considered here. In our notations, a GoE state X is characterized by having *P*_*X*_ = ∅, which happens only when there are two factors 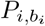 and 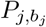 in 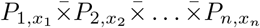 such that 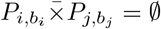. Given the commutative nature of 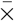, we only need to consider the *n*(*n* − 1)*/*2 possible products of pairs of 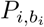 sets to determine the conditions under which X has no pre-image. A possible approach at finding the attractors of a boolean network consists in finding its GoE states first, and then following their evolution into their corresponding attractor states. An attractor is able to elude this search only when it coincides with its basin, e.g. a fixed point attractor with no other pre-image, or a pure loop of states with no branch attached to it.

### High copy number limit

In the last part of this section we will briefly discuss the cumulative effect of successive events of whole network duplication.

#### Preposition 3

In the very specific case of a network with zero thresholds (*B* = 0), the attractor landscape in *S*_21_ is a graph with exactly the same topology as the one in *S*_1_ describing the dynamics of *N*_1_. The two networks have then the *same attractors* under the mapping *X* ↔ (*X, X*), which is no longer just a set isomorphism, but a pseudo-digraph isomorphism between the attractor landscapes in *S*_1_ and *S*_21_.

**Proof:** Preposition 3 is an immediate consequence of equations 4 with *b*_*i*_ = 0.

In the reminder of this manuscript, whenever attractors of *N*_1_ and *N*_2_ are identified, the correspondence is implicitly established through the map described in Preposition 3.

The interest in this scenario comes from the fact that it represents the limiting case of multiple events of whole network duplication. We have seen that each duplication event reduces the thresholds in the network by a factor two. Therefore, after *m* events –with *m* large enough for the effective thresholds *θ*_*i*_*/*2^*m*^ to be smaller than the smallest differences in the terms 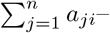 the thresholds are effectively zero, and remain so at each successive duplication. In the language of GRNs, the initial duplications determine a phenotypic shift, while further ones preserve the identity of the phenotypes and just reinforce the most stable ones by enlarging their relative basin of attraction after each duplication.

## Discussion

Motivated by its potential to describe whole gene duplication in systems dynamics models of gene regulation, we have analyzed the effect that network duplication has on the nature of the state space of dynamical Boolean networks with thresholds.

Since the seminal work of Ohno [14], gene and genome duplication have been ascribed to the main mechanisms responsible for the adaptive evolution of genetic systems [15]. Even if a much rarer evolutionary event than segmental duplication –involving from few nucleotides to several thousand kilobases– polyploidization –i.e. the duplication of the whole genome– has occurred repeatedly over geological time scales [16, 17]. Plants are among the organisms best know for having undergone multiple polyploidization events [18]. For example, three such events have occurred in the evolutinary history of *Arabidopsis thaliana* over a timescale of 150 million years [19]. Even in humans, ohnologs, i.e. paralogs retained from early duplications of the whole genome, constitute about 20 − 35% of the genome [20]. On non-evolutionary time, whole genome duplication has been observed as a stress response in heart and liver cells in both humans and mice [21]. Such spectacular events either lead to extinction [22] or re-diploidization, i.e. the generations of duplicated chromosomes differing from their templates in large segments. Not all chromosomes are retained, leading to the so called phenomenon of aneuploidy, i.e. chromosome loss [23].

Given the complexity, and the variety spanned by the individual scenarios, an exhaustive, mechanistic treatment of the phenomenon of whole genome duplication from a systems dynamics perspective is not just beyond the scope of this manuscript, but very likely premature. The more realistic objective of this manuscript has been to focus on the key role played by the symmetry of perfect polyploidy in shaping the attractor landscape of an idealized model for duplicated GRNs. In the real world, this symmetry is broken by diverging effects such as further mutations, fixation of duplicates and their selective preservation [24, 25]. Nonetheless, we believe this manuscript constitutes, beyond the systems dynamics techniques it highlights, a preliminary benchmark for organism-specific studies where the effect of divergence can be progressively superimposed to the unperturbed dynamics discussed here.

More specifically, we have shown that the symmetry of the regulatory network poses an enormous simplification in determining the phenotypes of the polyploid. As a result of that, solving the dynamics of the duplicated network is not harder than solving the dynamics of the network pre-duplication. We have also revisited the role that a reverse algorithm for finding the basins of attractions plays in these systems. While not computationally more advantageous than straightforward evolution in generic dynamical networks, in the case of a duplicated network its computational complexity is the same as that of the unduplicated network. This has been achieved by: 1) Reformulating the problem in terms of parametric sets whose size scales exponentially with the average connectivity of the network, and not anymore exponentially with the network size, and 2) By highlighting that the increased number of states of the duplicated network is entirely due to the acquisition of additional Garden of Eden states.

As long as asymmetries are ignored, these results allow to exactly solve the dynamics of networks resulting from an arbitrary number of duplication events. This poses the question of the effect that multiple duplication events have on the attractor states. We have show that –while the first duplication events can change the nature of the attractors– successive events can only affect and/or reinforce the size of the basins. We plan to further explore this latter result, especially in the light of the insights it could provide to plant systems biology, where such events are especially common [18].

## Footnotes

1 An exception is provided by self-referencing rules, e.g. when *x*_*i*_ and *x*_*n*+*i*_ receiving and integrated signal equal to their activating threshold means that they keep their values constant, and different, because their initial conditions were different.

